# Precision Targeting of Na-K-2Cl cotransporter (NKCC1) RNA In Vitro with RfxCas13d

**DOI:** 10.1101/2023.05.17.541102

**Authors:** Alfredo Sandoval, Bo Chen

**Author notes:** Correspondence (A.S), (B.C).

## Abstract

The Na-K-2Cl cotransporter (NKCC1) is considered an attractive drug target in the Central nervous system (CNS) for treating various CNS disorders. However, the specific role of NKCC1 in different types following injury within the CNS is not well understood due to its expression in multiple cell types. Additionally, there is a lack of a robust method for knocking down NKCC1 transcripts. In this study, we utilized Cas13 nucleases, a type of programmable RNA-targeting CRISPR enzyme, to effectively degrade NKCC1 mRNA in cultured cells. We developed a versatile pipeline for crRNA screening and validation in vitro and demonstrated the successful knockdown of NKCC1 using RfxCas13d. Our findings establish RfxCas13d as a powerful tool for targeting specific transcripts in vitro. By demonstrating the successful in vitro application of RfxCas13d-mediated NKCC1 RNA knockdown, we have laid the groundwork for future investigations into the therapeutic potential of NKCC1 modulation in CNS disorders.

## Introduction

Programmable RNA-targeting CRISPR-Cas systems constitute a diverse family of endonuclease enzymes that have served as valuable tools for DNA editing^1,2^ and more frequently, as promising platforms for transcriptome engineering. Among these systems, the Cas13 family of enzymes, such as RfxCas13d, have emerged as powerful tools for RNA knockdown^3-8^. Like its Cas9 counterpart, the Cas13 enzymes can be directed to target specific nucleic acid sequences via a programmable single stranded CRISPR RNA (crRNA). Several Cas13 subclasses have been identified and engineered for programmable RNA targeting^4,5,8^. Among these, RfxCas13d has emerged as a favored platform due to its small size – allowing it to fit in a single adeno-associated virus vector – and its high knockdown efficiency and on-target specificity^3^. High-fidelity versions of Cas13d have been recently engineered and discovered^7,8^, providing additional benefit over common knockdown techniques such as small hairpin RNAs (shRNAs). This combination of traits makes the RfxCas13d and its variants well suited for mechanistic studies that require low off-target effects with substantial knockdown efficiency.

In the context of CNS disorders, understanding the mechanism requires manipulating the genetic and pharmacological aspects of spinal cord and brain neurons following injury^9^. For instance, spinal neurons in spinal cord injury (SCI) often exhibit reversible and irreversible cell damage^10^, where unintended large-scale perturbations in transcriptome due to off target knockdown could increase the likelihood of apoptosis and necrosis. The NKCC1 plays a role in altering neuronal chloride regulation, impacting GABAergic signaling, and has been implicated in numerous CNS disorders. However, the specific role of NKCC1 in neurons following injury remains unclear due to its expression in various cell types within the CNS.

Given these observations and the benefits of Cas13d, we sought to investigate whether RfxCas13d could be used to target transcripts associated with spinal neuron silencing after SCI. In this study we demonstrate that RfxCas13d is a robust RNA targeting platform that can used to readily knockdown highly expressed NKCC1 in vitro. Furthermore, we establish a reliable workflow for knocking down disease associated mRNA transcripts, representing a crucial step towards future animal studies. This approach holds potential for the development of novel therapeutic strategies aimed at modulating ion channel expression for the treatment of various neurological disorders.

## Results

### Knockdown Strategy and Selection of Targeting Sequences

We sought to determine whether RfxCas13d could be used to knockdown mouse NKCC1, a gene of interest for functional recovery after spinal cord injury. To accomplish this, we first devised a strategy to select the most effective CRISPR RNAs (crRNAs) for targeting the mouse NKCC1 transcript. We designed 5 crRNAs with 23-nucleotide (nt) protospacers targeting coding exons of the NKCC1 transcript. Our selection was based on predicted knockdown efficiency and minimal off-target effects using the web-based tool, Cas13design (Figure 1). To identify crRNAs that could most effectively knockdown NKCC1, we designed an NKCC1 reporter expression vector that harbors the NKCC1 sequence after an mCherry reading frame, to minimize localization artifacts and potential toxicity from NKCC1 overexpression.

**Figure 1.**
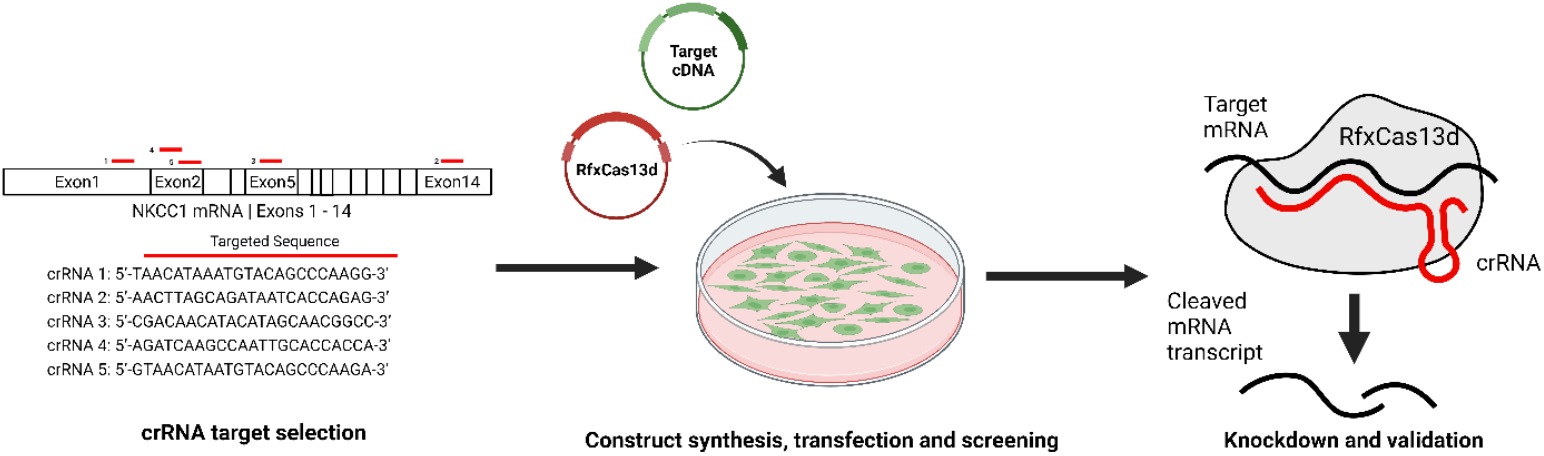
Schematic of Cas13d knockdown strategy. CRISPR RNAs (crRNAs) targeting the mouse NKCC1 transcript were chosen based on predicted knock down efficiency and off-targets (Cas13design). Selected targeting sequences were synthesized and co-transfected with an NKCC1-mCherry overexpression vector into 293FT cells and analyzed for knockdown efficiency.

### Cas13d-Mediated Knockdown of NKCC1 RNA

To assess the effectiveness of RfxCas13d in mediating the knockdown of NKCC1 RNA, we transfected RfxCas13d-crRNA expression constructs with the NKCC1-mCherry reporter into 293FT cells to visually determine the extent of knockdown in vitro. Representative images from these assessments are depicted in Figure 2. The samples included (A) mCherry-NKCC1 alone, (B) RfxCas13d with a control guide, (C) NKCC1 Guide 3, (D) NKCC1 Guide 4, and (E) RfxCas13d-2A-EGFP alone. These results indicated that RfxCas13d can effectively harness specific guide RNAs to knockdown NKCC1 mRNA.

**Figure 2:**
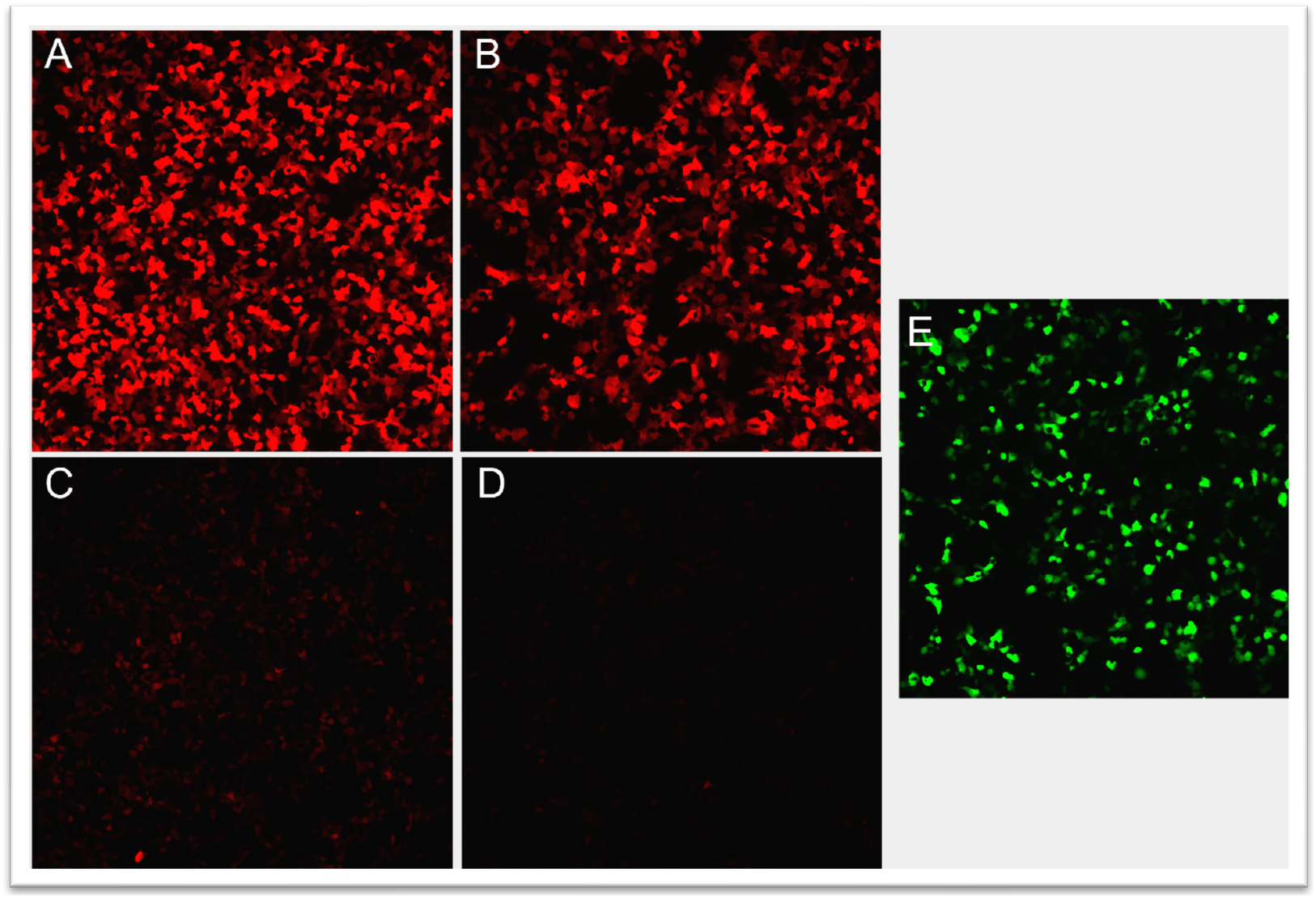
Cas13d-Mediated Knockdown of NKCC1 RNA. Representative images of RfxCas13d mediated knockdown in 293FT cells. (A) mCherry-NKCC1 alone (B) RfxCas13d with control guide, (C) NKCC1 Guide 3, (D) NKCC1 Guide 4, (E) RfxCas13d-2A-EGFP alone

### Quantitative Analysis of Cas13d-Mediated Knockdown

To quantify our visual assessments, we performed a quantitative analysis of the mCherry Mean Fluorescence intensity (MFI) in HEK293FT cells transfected with the NKCC1 reporter and RfxCas13d-crRNA constructs. This analysis was conducted 24 hours after transfection. Our results, shown in Figure 3, revealed a significant reduction in MFI in cells transfected with NKCC1 Guide 3 and Guide 4, compared to the LacZ and reporter only control, indicating successful NKCC1 knockdown (One-way ANOVA, N = 3 replicates per condition, Error Bars indicate SEM, ***P=0.0001, ****P<0.0001). Meanwhile, no significant difference was observed with the control guide.

**Figure 3:**
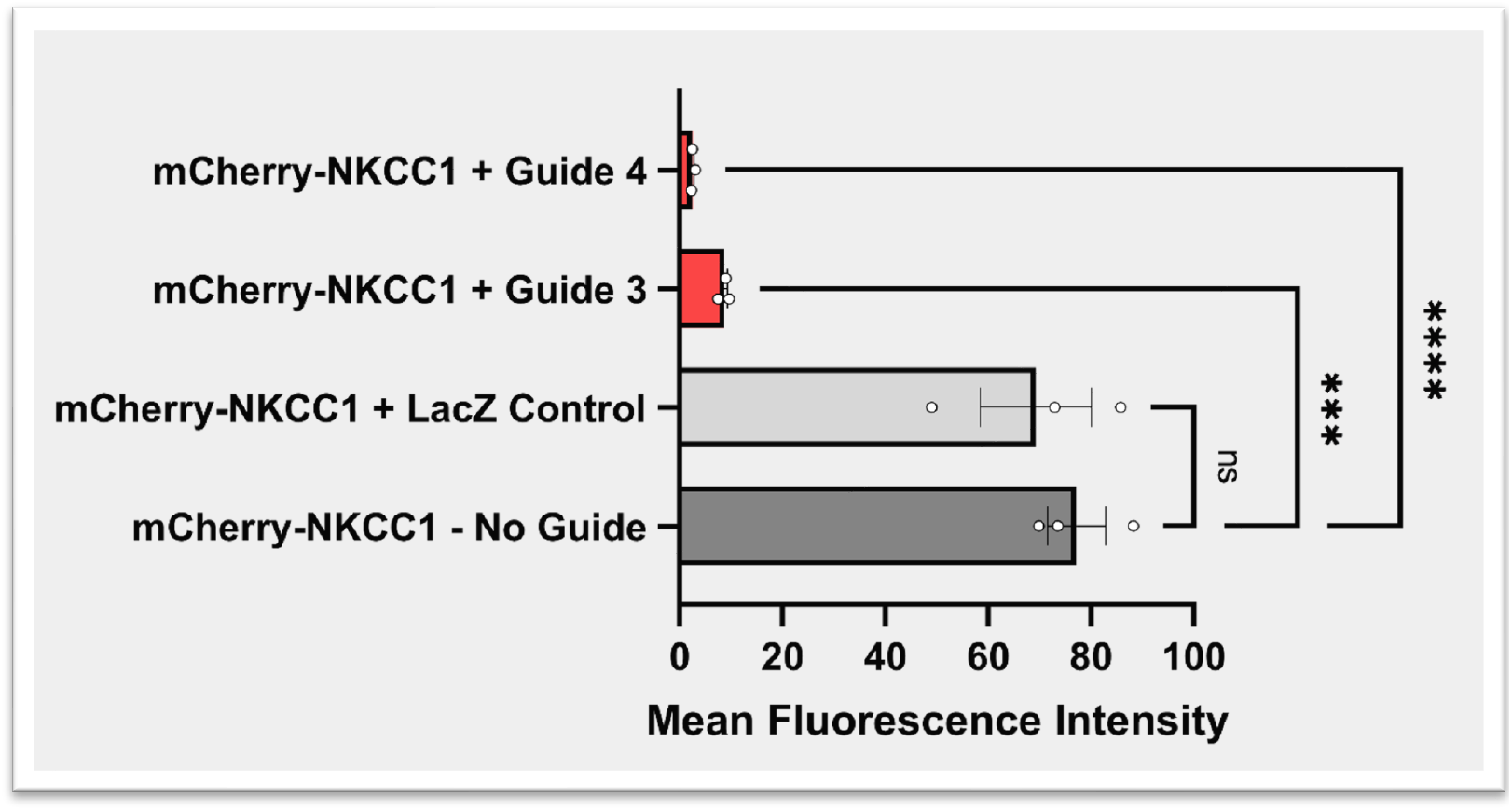
Quantitative analysis of Cas13d-Mediated Knockdown of NKCC1 RNA from Figure 2. mCherry Mean Fluorescence intensity (MFI) in HEK293FT cells transfected with NKCC1 reporter and RfxCas13d-crRNA construct. All comparisons (One-way ANOVA) are compared to reporter only control 24 hours after transfection. N = 3 replicates per condition. Error Bars indicate SEM. ns = not significant, ***P=0.0001, ****P<0.0001

In conclusion, our results provide evidence that RfxCas13d can be successfully harnessed for the knockdown of NKCC1 RNA in vitro. This proof-of-principle study serves as a key step towards validating the feasibility and efficacy of RfxCas13d in modulating gene expression in more complex in vivo models.

## Conclusion

In this study, we have successfully demonstrated the potential of RfxCas13d-mediated knockdown for precise and targeted gene regulation in vitro. We have utilized RfxCas13d to specifically target and knockdown NKCC1 RNA, a crucial component of the Cation Cl-Cotransporters (CCCs) family. NKCC1 is implicated in various physiological and pathological processes including neuronal development, cell volume regulation, and neurological disorders, especially spinal cord injury (SCI). Dysregulation of NKCC1 following SCI can result in detrimental cellular changes, such as neuronal swelling and hyperexcitability, which contribute to the progression of the injury and the associated functional deficits.

By demonstrating the successful in vitro application of RfxCas13d-mediated NKCC1 RNA knockdown, we have laid the groundwork for future investigations into the therapeutic potential of NKCC1 modulation in SCI models. The potential to modulate NKCC1 expression using RfxCas13d may provide novel insights into the role of NKCC1 in SCI pathology and could open new avenues for the development of targeted therapeutic strategies.

Future work will extend the use of RfxCas13d-mediated NKCC1 modulation to in vivo models of SCI, with the aim of elucidating the biological consequences of NKCC1 expression and its therapeutic impact in the context of SCI. We anticipate that these investigations will further our understanding of the role of NKCC1 and other CCCs in SCI pathology and may contribute to the development of novel therapeutic interventions aimed at modulating ion transport and cell volume regulation in neurons.

## Methods

### Plasmid Synthesis

All plasmids used in this study were synthesized by Genscript. The pAAV-EFS-Cas13d-2A-EGFP plasmid was derived from pXR001 (Addgene ID: 109049). The CAG-mCherry-NKCC1 plasmid was based on pcDNA3.1(+)-C-eGFP. All constructs were confirmed by whole plasmid sequencing (plasmidsaurus).

### Targeting Sequence Design

Targeting sequences for the Cas13d-mediated knockdown were designed using Cas13design, a web-based tool available at https://cas13design.nygenome.org/. Targeting sequences were carefully selected based on their predicted specificity and efficiency to ensure accurate and effective knockdown of mNKCC1 RNA.

### Cell Culture and Transfections

293FT cells (Invitrogen) were cultured according to the manufacturer’s instructions in DMEM high glucose (Corning) with 10% FBS (Gibco) and 1% penicillin-streptomycin (Gibco). Twenty-four hours before DNA transfection, cells were seeded at a density of 150,000 cells per well in a 24-well cell culture plate.

Equal amounts of plasmid were transfected into the cells using Lipofectamine 3000 (Invitrogen) following the manufacturer’s instructions. The transfections were performed with the same technique for all experiments to maintain consistency and minimize technical variability.

### Imaging

Cells were imaged on an Olympus IX81 microscope using the appropriate filters at 10x magnification. Imaging sessions were conducted 12 and 24 hours after transfection to monitor cellular changes over time. To ensure consistency in image acquisition, the same exposure settings were applied to all samples. Cells were imaged in FluoroBright media (Gibco) to provide optimal conditions for fluorescence visualization and minimize background fluorescence.

### Statistical Analysis

Statistical analysis was performed using GraphPad Prism 8. Mean Fluorescence Intensity of each replicate was determined with ImageJ (NIH) and were compared using one-way analysis of variance (ANOVA).

## Funding

This study was supported by grants from University of Texas Medical Branch (UTMB) startup funds (B.C.); Mission Connect, TIRR Foundation (B.C.); UT system star award (B.C.); Wings for Life Foundation Research grant (B.C.)

## Notes

### Competing Interest Statement

The authors have declared no competing interest.

